# Acetyl-CoA carboxylase 1-dependent lipogenesis drives breast cancer progression

**DOI:** 10.1101/2023.07.20.549828

**Authors:** Keely Tan, Thomas Owen, Holly P. McEwen, Peter Simpson, Andrew J. Hoy, David E. James, Anthony S. Don, Matthew J. Naylor

## Abstract

Dysregulation of cellular energetics, including lipid synthesis mediated through de novo lipogenesis, is a feature of many cancers. Here we report that acetyl-coenzyme A carboxylase (ACC) 1, the rate-limiting enzyme of de novo lipogenesis, is a key regulator of breast cancer progression and cancer cell phenotype. Mammary epithelial-specific deletion of ACC1 impaired tumour progression and decreased cancer cell proliferation in the PyMT model of breast cancer *in vivo*. ACC1 knockout in human breast cancer cell lines resulted in decreased cell number and altered cell and membrane morphology. Lipidomic profiling demonstrated reduced levels of acyl-carnitines (CARs) and several phospholipid (PL) classes, whilst also shifting the lipid profiles to exhibit more elongated and less saturated lipids in ACC1 knockout breast cancer cells. Palmitate rescue of ACC1 deletion phenotypes demonstrated a critical role for ACC1 driven de novo lipogenesis in breast cancer cell function. Analysis of human breast tumour-microarrays identified strong ACC1 expression at all breast cancer stages, grade and metastasis, compared to normal adjacent tissue. Together our data demonstrate a novel role for ACC1 in breast cancer progression and cancer cell function, mediated through its lipogenic role, that together with its expression profile, identify ACC1 as a potential therapeutic target in breast cancer.

**Statement of significance:** This study investigates the impact of ACC1 deletion in breast cancer progression, revealing the importance of ACC1-derived lipids in breast cancer cell phenotypes and identifies ACC1 as a potential novel therapeutic target.

## Introduction

Breast cancer is a complex and heterogeneous disease that accounted for 25% of cancer diagnoses and 16% of cancer-related deaths in women in 2020 alone, making it the most commonly diagnosed and lethal cancer of women [1]. With reprogramming of cellular metabolism during tumourigenesis recognised as a hallmark of cancer [2], lipids are emerging as a metabolic source during cancer progression. Lipids can be derived from both exogenous and endogenous sources [3], but there is increasing evidence that cancers are reliant on *de novo* sources irrespective of circulating lipid levels [4, 5]. The process of *de novo* lipogenesis (DNL) is governed by the acetyl coenzyme-A (CoA) carboxylases (ACCs), the rate limiting enzymes of the *de novo* lipid synthesis pathways. Previous cancer research has predominantly focused on the ACC1 isoform, given its known role in lipid biosynthesis, and the importance of lipids as a macromolecule for tumour biomass generation [6].

A potential role for ACC1 in breast cancer has been inferred by the link between ACC1 and the tumour suppressor breast cancer susceptibility protein (BRCA) 1 [7] and human epidermal growth factor receptor (HER) 2 overexpression [8]. *In vitro*, ACC1 has been shown to influence breast cancer cell proliferation, survival and stem cell renewal [9, 10]. The mechanism of how this occurs is potentially due its effect on lipids, particularly phospholipid characteristics, influencing cell mechanics involved in cell proliferation and motility [9]. In contrast, Garcia and colleagues indicated that ACC1-dependent protein acetylation contributes to breast cancer metastasis and recurrence [11], indicating that the role of ACC1 driven DNL remains to be elucidated. While *in vitro* studies are beneficial for preliminary and mechanistic research, they fail to replicate the conditions of cells in an organism [12]. Of the limited available studies, most used non-specific ACC inhibitors to investigate the role of ACC1 in breast cancer, which is unable to distinguish between the individual contribution of the two different ACC isoforms. Thus, the aim of this study was to specifically investigate the role of ACC1-derived lipids in breast cancer development and progression, both *in vitro* and *in vivo*, using ACC1 specific genetic knockout models.

This present study reports for the first time, an *in vivo* role for ACC1 in mammary tumorigenesis, particularly through its role in driving cancer cell proliferation. This is further confirmed in our *in vitro* studies, where we demonstrate genetic deletion of ACC1 in breast cancer cell lines decreased live cancer stem/progenitor cells, due to altered cell morphology resulting from lipid dysregulation. Subsequent lipidomic studies highlight a critical impact of ACC1 deletion on the levels of acylcarnitines (CARs) and several phospholipid (PL) classes. In addition to an overall reduction in these lipids, ACC1 deletion also shifted lipid profiles to exhibit more elongated and less saturated lipids. Addition of palmitic acid rescued the cell morphology changes and decreased cell number caused by ACC1 deletion, confirming the contribution of ACC1-derived lipids to breast cancer cell phenotypes. To explore the clinical relevance of ACC1, this study highlights the importance of the enzyme in the initiation of human breast cancer, given its upregulation in malignant tissue when compared to normal adjacent tissue.

## Methods

### Longitudinal study

All animal experimentation was carried out in accordance with University of Sydney Animal Ethics Approval 2017/1204. ACC1 floxed mice [13] were crossed with Blg promoter driven Cre recombinase (BlgCre) transgenic mice [14], resulting in mammary-specific ACC1 KO mice. These mice were then crossed with the polyomavirus middle T antigen (PyMT) breast cancer mouse model [15, 16] which were under the control of mouse mammary tumour virus LTR (MMTV LTR) to create female wildtype (WT) (ACC1^f/f^BlgCre^+/+^PyMT^tg/+^) or ACC1 mammary-specific epithelial KO (ACC1^f/f^BlgCre^tg/+^PyMT^tg/+^) mice on the PyMT driven breast cancer tumour model. Mice were euthanised when the tumour mass reached the ethical end point of 10% of total body weight.

### Cross-sectional mouse tissue collection

Abdominal mammary glands and lungs were collected from female ACC1^f/f^BlgCre^tg/+^PyMT^tg/+^ and ACC1^f/f^BlgCre^+/+^PyMT^tg/+^ mice at 8 and 11 weeks.

### Mammary gland mounting and gland imaging

Abdominal mammary glands were fixed overnight in 10% neutral buffered formalin (NBF), and then defatted the following day using three changes of 100% acetone every 2 hours. Following this, glands were incubated in 70% ethanol for 1 hour before staining overnight in carmine alum (0.2% carmine, 0.5% aluminium sulphate). The gland was then dehydrated in a series of ethanol washes (70-100%) for 2 hours each before washing in xylene for 1 hour. Finally, glands were stored in methyl salicylate (Sigma-Aldrich) for imaging. Images were taken using the Leica M165FC microscope. All mammary gland slides were imaged at 4 different magnifications; 0.73X, 1.6X, 2.5X and 5X. Following imaging, glands were rehydrated through a series of overnight ethanol baths (100%, 95% and 70%), and stored in 70% ethanol for embedding and sectioning. Sections were then paraffin embedded and sectioned for further analysis.

### Immunohistochemistry

Histological staining was performed on paraffin sections of mammary glands and human tumour microarray (TMA) slides (BR2085d and BR10010b, US Biomax). Slides were dewaxed by pre-warming, followed by xylene washes and hydrated in a series of ethanol washes (100%-70%). Antigens were retrieved using Target Retrieval Solution, High pH (Dako Omnis) (Agilent) in a heated water bath. Endogenous peroxidases were deactivated using 3% H_2_O_2_, and then primed with background sniper (MACH 1 Universal HRP-Polymer Kit, Biocare Medical). Following this, sections were incubated with ACC1 (ab72046, 1:1000), Ki67 (MA5-14520, 1:100) or anti-rabbit IgG (7074, 1:20000) at room temperature, followed by incubating with a secondary antibody. To visualise, sections were treated with DAB substrate, counterstained with haematoxylin and mounted with Safety Mount No. 4 (Fronine). For mammary glands, slides were imaged and positive immunostaining was calculated from 4 randomly selected images using ImmunoRatio Image J plug-in. Cores from human TMAs were imaged and scored based on a scoring index created from samples from the slide.

### Haematoxylin and eosin (H&E) staining

Following dewaxing steps above, sections were stained in haematoxylin, and washed until water was colourless. Slides were then dipped in 1% acid alcohol for 10 seconds and continued with Scott’s Bluing reagent. Dehydration was initiated through a 70% and 95% ethanol wash, for 1 minute each, before dipping in eosin Y. Sections were dehydrated in a series of ethanol washes (95%, 95%, 100% and 100%). Slides were then mounted with Safety Mount No. 4 (Fronine). Slides were scanned using the ZEISS Axio Scan.Z1 slide scanner and stitched to form a complete virtual image. Percentage of haematoxylin staining was calculated by separating haematoxylin and eosin into separate colour channels using the Colour Deconvolution Image J plug-in. The percentage of haematoxylin was then calculated as a percentage of the total mammary gland area.

### Cell culture

MDA-MB-231 and MCF7 human breast cancer cell lines purchased from ATCC (USA) were tested for mycoplasma contamination, using the MycoAlert™ PLUS Mycoplasma Detection Kit (Lonza). MDA-MB-231 cells were maintained in RPMI 1640 media supplemented with 10% FBS, 2mM L-glutamine (Gibco) and 100U/mL Penicillin-Streptomycin (Gibco). MCF7 and cells were also grown in the above media, with the addition of 10mM HEPES. Cells were kept at constant conditions of 37°C and 5% CO_2_ in an incubator. All cell experiments were performed in RPMI or DMEM media supplemented with 5% charcoal stripped FBS (cFBS) (Gibco) as a method of removing substantial amounts of lipophilic materials from the media [17, 18], with all other appropriate components present.

### CRISPR ACC1 knockout cell generation

CRISPR targeting sequences for ACC1 (F: GGCTTGCACCTAGTAAAGCA, R: TGCTTTACTAGGTGCAAGCC) obtained from Sigma were cloned into the pSpCas9(BB)-2A-GFP bacterial plasmid sourced from Addgene (PX458; RRID: Addgene_48138) following the protocol as defined by the Zhang lab [19] and sequenced for correct oligonucleotide insertion. The plasmid was then transfected into MDA-MB-231 and MCF7 cells for 72 hours before cells were sorted based on GFP expression using flow cytometry into 96 well plates to obtain single colonies. Media was supplemented every 48-72 hours and expanded for freezing and protein extraction. Knockout of protein was validated by western blotting. MDA-MB-231 and MCF7 cells transfected with an empty plasmid and selected for GFP expression were used as controls in subsequent CRISPR experiments, unless otherwise specified. Cells were imaged 72 hours post treatment in 5% cFBS to visualise morphological changes and after 168 hours, cells were trypsinised and cell viability was calculated by trypan blue exclusion with a haemocytometer.

### Mammosphere formation assay

ACC1 KO cells were strained using 5mL polystyrene round-bottom tubes (BD Falcon, #352235) and cultured in ultra-low attachment 6-well plates. Mammospheres were defined as spherical growths ≥ 50μm in diameter and did not resemble loosely clustered cells. After 96 hours, fresh conditioned media was added to each well. Images of mammospheres were taken 168 hours after plating using the ZEISS Axio Vert.A1 microscope and Zen 2012 (Blue Edition) software (ZEISS). After 168 hours, mammospheres were collected and dissociated into single cells, and live cells counted by trypan blue exclusion with a haemocytometer. Mammospheres were analysed for size and number using the Photoshop (Adobe) ruler tool, by creating a measurement scale preset based on the scale bar of the microscope captured images and used for all subsequent images.

### Palmitate rescue

Control and ACC1 KO cells were plated overnight and supplemented with 5% cFBS media and palmitic acid (palmitate) the following day. MDA-MB-231 cells were treated with 5µM palmitic acid and MCF7 cells were treated with 25µM palmitic acid. After 72 hours, wells were imaged using phase-contrast microscopy on the ZEISS Axio Vert.A1 microscope to identify the effects on morphology. After 168 hours, cells were trypsinised and cell viability was calculated by trypan blue exclusion and a haemocytometer.

### Immunofluorescence and imaging

ACC1 CRISPR KO cells were cultured onto 15mm round coverslips (Menzel Gläser). Immunofluorescence steps were performed at RT unless specified. Seventy-two hours later, cells were fixed in 4% PFA for 10 minutes, before washing with PBS tween (0.1%) and blocked (5% goat serum, 0.1% Triton X100, 0.1% bovine serum albumin diluted in PBS) for 2 hours. Cells were incubated with anti ß-tubulin antibody (1:500, Cell Signaling Technology #2146, RRID: AB_2210545) at 4°C overnight. The following day, cells were washed PBS tween before incubating with anti-rabbit IgG AlexaFluor^®^ 647 conjugate (Cell Signaling Technology, #3452, RRID: AB_10695811) for 2 hours and washed again in PBS tween. Cells were then incubated with DyLight™ 594 Phalloidin (Cell Signaling Technology, #12877) for 15 minutes and rinsed with PBS before incubating with DAPI for 7 minutes. After staining, cells were washed in PBS tween and mounted using fluorescent mounting medium (Agilent Dako, #S3023) for imaging. Images were obtained using the ZEISS Axio Scope.A1 microscope and Zen 2012 (Blue Edition) software (Zeiss) at identical settings between images, and analysis was performed using ImageJ version 2.0. Intensity was quantified using thresholding, and corrected for background fluorescence to calculate the corrected total cell fluorescence (CTCF) as previously described by Hammond [20]. To calculate cell area, thresholding was used to calculate the total cell area in each image, divided by the number of cells, determined by counting the number of nuclei using DAPI. Membrane ruffles were counted manually using ImageJ software.

### Lipid extraction

MDA-MB-231 and MCF7 ACC1 KO cells were treated for 72 hours with 5% cFBS media, then washed and trypsinised for pelleting. Lipids were then extracted following the two-phase procedure using methyl-*tert*-butyl-ether (MTBE)/methanol/water (10:3:2.5 v/v/v) [21]. Briefly, on the day of extraction, 200µL of high-performance liquid chromatography (HPLC) grade methanol containing 0.01% 3,5-di-tert-4-butylhydroxyltoluene (BHT) was added to each sample and spiked with an internal standard mixture (Supplementary Table S1) for post-acquisition normalisation. Three rounds of lipid extraction were performed using MTBE and methanol (containing 0.01% BHT), and then sonicated for 30 mins in an ice-cold water bath (Thermoline Scientific, Australia). For each round, phase separation was induced through the addition of milliQ water and completed by vortex and centrifugation of samples at 2000 x *g* for 5 minutes. The upper organic MTBE layer removed from each round of lipid extraction was combined and dried in a Savant SC210 SpeedVac (ThermoScientific). After samples were dried down, lipids were reconstituted in 200µL of methanol (containing 0.01% BHT), vortexed and centrifuged at 2000 x *g* for 10 minutes to pellet insoluble material. Extraction was completed by removing 150µL of the soluble material into a HPLC vial for storage.

### Lipid profiling and quantification

Lipidomic profiling was performed using untargeted liquid chromatography-tandem mass spectrometry (LC-MS/MS) on a QExactive HF-X mass spectrometer with a heated electrospray ionisation probe and a Vanquish HPLC (ThermoFisher Scientific) [22]. Extracts were resolved on a 2.1 × 100 mm Waters Acquity C18 UPLC column (1.7 µm pore size), using a binary gradient in which mobile phase A consisted of acetonitrile:water (60:40), 10 mM ammonium formate, 0.1% formic acid; and mobile phase B was isopropanol:acetonitrile (90:10), 10 mM ammonium formate, 0.1% formic acid. The flow rate was 280µL/min and the column oven was 45°C. Data was acquired in both positive and negative mode using full scan/data-dependent MS2 (full scan resolution 60,000 FWHM, scan range 220–1600 m/z). The 10 most abundant ions in each cycle were subjected to MS2 using an isolation window of 1.1 m/z, collision energy 30 eV, resolution 15,000, maximum integration time 35ms and dynamic exclusion window 8 s. An exclusion list of background ions was used based on a solvent blank. LipidSearch v4.2.21 software (ThermoFisher Scientific) was used for chromatogram alignment, peak identification and integration. Each lipid peak was manually verified for accurate precursor *m/z* (mass accuracy 5 ppm) and characteristic fragment ions (mass accuracy 8 ppm), as well as column elution time. Data were then exported to an Excel spreadsheet for normalisation of each lipid to its class-specific internal standard, and lipid concentrations were calculated relative to the internal standard. The top fifteen most abundant lipid species in each class were then used for further analysis. Data was either log_2_ transformed or normalised as a percentage of total lipid pool for each lipid class for graphical purposes. MetaboAnalyst 4.0 was used for generation of heat maps and volcano plots.

### Western blot

Protein from cells was extracted RIPA lysis buffer. Lysates were chilled on ice for 10 minutes, before centrifuging for 20 minutes (17, 000 x *g* at 4°C). Total protein isolates were quantified by the Direct Detect^®^ FTIR spectrometer. Samples (20µg) were resolved on 4-15% Mini-PROTEAN^®^ TGX^™^ Precast Protein Gels (BioRad) in running buffer (25mM Tris, 192mM glycine and 0.1% (w/v) SDS, pH 8.3) using the Mini-PROTEAN^®^ electrophoresis system (BioRad), with Precision Plus Protein™ Dual Color (BioRad) as the molecular marker. The cassette was created according to the Trans-Blot Turbo Blotting System (BioRad) and transferred onto a polyvinylidene fluoride (PVDF) membrane using the Trans-Blot Turbo. The membrane was blocked with 5% skim milk for 1 hour at RT before immunoblotting with antibodies against ACC1 (#ab72046, Abcam, 1:2000), ACC2 (#HPA006554, Sigma-Aldrich, 1:500), CPT1A (#ab128569, Abcam, 1:1000), GAPDH (#ab8245, Abcam, 1:2000), vinculin (#ab219649, Abcam, 1:2000) at 4°C overnight. The following day, membranes were washed 4 times using 0.2% TBS tween before probing with the secondary antibody (anti-mouse (#7076; AB_330924) (1:10000) or anti-rabbit (#7074; RRID: AB_2099233) (1:2000) IgG, HRP-linked antibody; Cell Signaling Technology) for 2 hours at RT. To visualise bands, membranes were washed in 4 x 5 minutes in 0.2% TBS tween before incubating in Clarity^™^ Western ECL Substrate (BioRad) for 5 minutes, and imaging using the BioRad ChemiDoc Touch. Densitometry was performed using ImageJ software and normalised in Excel.

### Statistical Analysis

All statistical analyses were performed using the GraphPad Prism 8 software (San Diego, USA). Graphs were also generated using the GraphPad Prism 8 software (San Diego, USA). Data are represented as mean ± standard error (SEM) or mean ± standard deviation (SD) and differences between groups were assessed using appropriate statistical tests, as detailed in figure legends. Results were considered statistically significant at p≤0.05.

### Data Availability

All data generated in this study are available upon request from the corresponding author.

## Results

### ACC1 deletion *in vivo* decreases tumour growth and progression

To investigate the effects of ACC1 deletion on mammary tumorigenesis, ACC1 floxed mice [13] were crossed with BlgCre recombinase transgenic mice [14] and subsequently crossed to PyMT^tg/+^ mice to create control (ACC1^f/f^BlgCre^+/+^PyMT^tg/+^; WT) or ACC1 mammary-specific epithelial KO (ACC1^f/f^BlgCre^tg/+^PyMT^tg/+^; KO) mice on the PyMT driven mammary cancer tumour model. PyMT transgenic mice develop luminal mammary cancers comparable with human ductal breast cancers of the luminal B subtype [15]. We first confirmed that ACC1 deletion was mammary gland and epithelial cell specific in these mice (Figure 1A). A longitudinal analysis demonstrated that ACC1 deletion increased median tumour-free period from 70 days in WT mice to 104 days in KO mice (p≤0.0001) (Figure 1B) and overall survival increased from 110 days in WT mice to 158 days in KO mice (p≤0.0001) (Figure 1C). To identify which stage of mammary tumour progression was influenced by ACC1, we performed a cross-sectional analysis at two time points. At both 8 (advanced pre-malignant lesion, p=0.0129) and 11 (early to late carcinoma, p=0.0173) weeks, the number of hyperplastic lesions was decreased (Figure 1D). This was accompanied with a 20.9% decrease at 8 weeks (p≤0.0001) and 9.4% at 11 weeks (p=0.0233) in proliferation, as measured by the percentage of Ki67 positive cells (Figure 1E). These findings suggest that ACC1 influences mammary cancer progression by driving epithelial cell proliferation in the luminal B PyMT breast cancer model.

**Figure 1.**
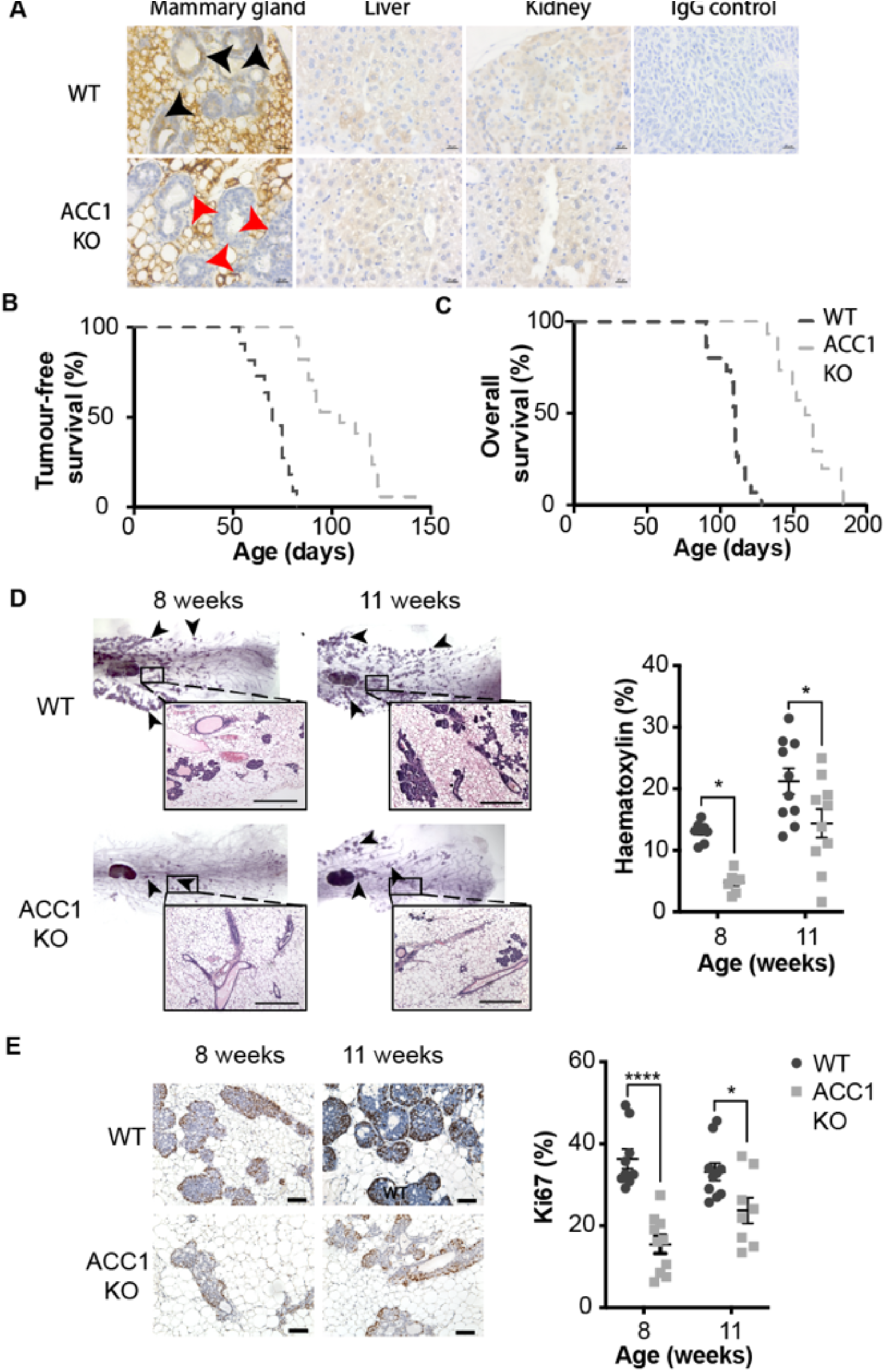
ACC1 deletion *in vivo* decreases tumour growth and progression. **(A)** ACC1 staining in 11-week-old mammary glands, liver and kidney from WT and ACC1 KO mice. Black arrows indicate ACC1 epithelial staining and red arrows indicate the absence of ACC1 in mammary gland tissue. Scale bar = 20μm. **(B)** Tumour-free (hazard ratio=0.17) and **(C)** overall survival (hazard ratio=0.18) of WT mice compared to ACC1 KO mice. **(D)** Whole mammary glands from 8-week and 11-week-old WT and ACC1 KO mice. Glands were stained with haematoxylin & eosin for microscopic analysis and quantified as a percentage of the whole gland area. **(E)** Immunohistochemical images of Ki67 staining in 8-week and 11-week-old WT and ACC1 KO mice. Ki67 positive nuclear staining of mammary glands was quantified as a % of total nuclear area. Scale bar = 100μm. Data is represented as the mean ± SEM (n=7-10 independent experiments, depending on group). Statistical significance for the survival curve was determined by a log-rank (Mantel-Cox) test and haematoxylin and Ki67 (%) was determined by a two-way ANOVA with Šídák’s multiple comparisons post-hoc test. *P≤0.05, ****P≤0.0001.

### ACC1 deletion *in vitro* decreases number of live cancer cells and alters cancer cell morphology

We next investigated the effects of ACC1 knockout in two different breast cancer cell lines, MDA-MB-231 and MCF7, to confirm and extend previous *in vitro* studies [9, 10, 23, 24]. The MDA-MB-231 cell line represents the triple-negative, poor prognosis, breast cancer subtype, while the MCF7 cell line represents luminal A, good prognosis, breast cancers [25]. ACC1 was deleted in the two cell lines using CRISPR-Cas9 technology and confirmed by western blot analysis (Figure 2A). Knockout of ACC1 decreased live cell number by ∼60% in both MDA-MB-231 (KO 1: p=0.0039; KO 2: p=0.0081) and MCF7 (KO 1: p=0.0054; KO 2: 0.0019) (Figure 2B) cell lines cultured in cFBS. We next used a low-adhesion mammosphere assay to assess cancer progenitor cell function and survival [26], following ACC1 deletion. ACC1 deletion resulted in ∼50-60% decreased live MDA-MB-231 (KO 1: P=0.0070; KO 2: p=0.0444) and MCF7 (KO 1: p=0.0284; KO 2: 0.0018) (Figure 2C) cells in the low-adhesion mammosphere assay.

**Figure 2.**
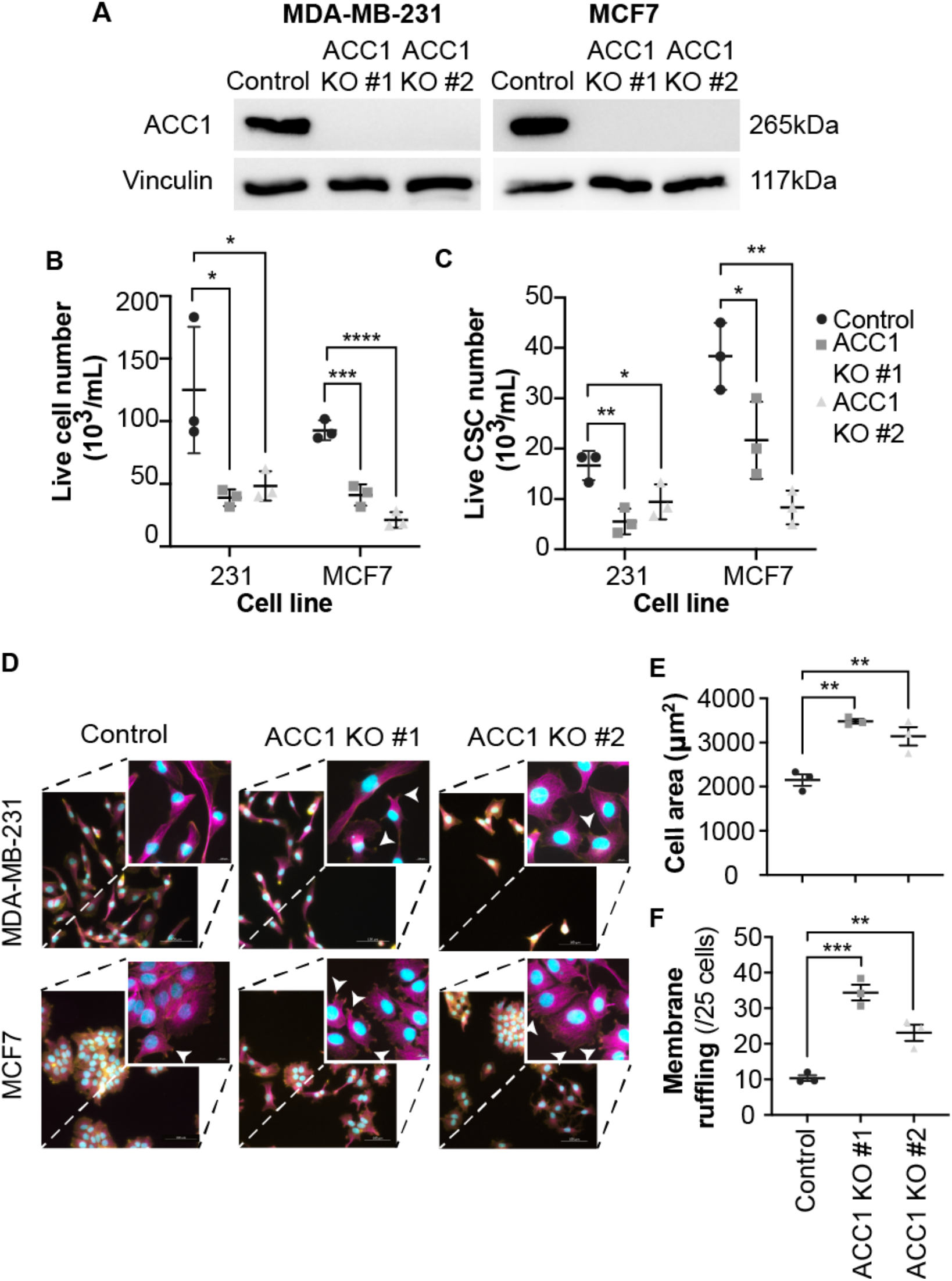
Deletion of ACC1 decreases cell number and alters cell morphology. **(A)** Western blot images confirming the CRISPR-Cas 9 mediated knockout of ACC1 in MDA-MB-231 and MCF7 cells. **(B)** Live cell count of MDA-MB-231 and MCF7 ACC1 knockout cells. **(C)** Number of live cells counted of control and ACC1 KO MDA-MB-231 and MCF7 CSCs. **(D)** Representative images of control and ACC1 KO MDA-MB-231 and MCF7 cells that were stained with fluorescent ß-tubulin (pink) and F-actin (yellow). Scale bar = 100μm. Representative images showing membrane ruffling (indicated by white arrows) in control and ACC1 KO MDA-MB-231 and MCF7 cells (panels within boxes). **(E)** Size of control and ACC1 KO MCF7 cells were quantified using ImageJ. **(F)** Quantification of membrane ruffling using ImageJ software in control and ACC1 KO MCF7 cells. Data is represented as the mean ± SEM (n=3 independent experiments). Statistical significance was determined by a one or two-way ANOVA with Dunnett’s multiple comparisons post-hoc test. *P≤0.05, **P≤0.01, ***P≤0.001, ****P≤0.0001.

We next investigated the effect of ACC1 deletion on MCF7 and MDA-MB-231 cellular morphology. MCF7 ACC1 KO cells exhibited altered morphology, such that they lost their ability to form tight clusters and appeared to have longer and thinner cytoplasmic processes (Figure 2D). This was accompanied with an increase in cell area (KO 1: p=0.0012; KO 2: p=0.0053) (Figure 2E) and increased membrane ruffling (KO 1: p=0.0002; KO 2: p=0.0057) (Figure 2F). In contrast there was no significant change in MDA-MB-231 cell morphology, membrane ruffling or cell size (Figure 2D, Supplementary Figure S2A-B), indicating inherent differences in the role of ACC1 in different breast cancer subtypes.

### Lipidomic analysis reveals alterations and re-distribution of the lipid profile following ACC1 deletion

To explore the mechanisms behind ACC1 regulation of breast cancer cell survival and maintaining cell morphology, liquid chromatography-tandem mass spectrometry was performed on lipids extracted from MDA-MB-231 and MCF7 cells cultured in low lipid media conditions. Deletion of ACC1 decreased various lipid species in MDA-MB-231 cells, in particular, acylcarnitine (CAR), phosphatidylcholine (PC), phosphatidylethanolamine (PE) and phosphatidylserine (PS) lipid classes (Figure 3A, Supplementary Figure 3A). Similarly, several lipid species were reduced in MCF7 cells following ACC1 deletion, and this was most prominent in the CAR, PC and PS classes (Figure 3B, Supplementary Figure S4A). Whilst the reduction of certain lipid species may be expected given the role of ACC1, there was also a shift in characteristics of lipid profiles. For example, the effect of lipid acyl-chain length has been reported in a range of cancers [27, 28]. Using the definition that shorter chain lipids equal a sum of the two-acyl chains ≤ 35 carbons and longer chains are ≥ 36 carbons, we calculated the elongation ratio [27], and observed increased longer acyl chains in PE and PS lipids in ACC1 KO MDA-MB-231 cells (Figure 3C, Supplementary Figure S5). Likewise, similar patterns were observed in CARs. For CARs, longer chains were defined as ≥ 18 carbons and shorter chains defined as ≤17 carbons lipids. Acyl chain saturation is another influential factor in cancer biology [3]. For CARs, longer chains were defined as ≥ 18 carbons ad shorter chains defined as ≤17 carbons lipids. The levels of unsaturated phospholipids (Figure 3D) and CARs (Figure 3E) were increased in MDA-MB-231 and MCF7 cells following ACC1 deletion. Overall, there was an increase in PUFAs in PC, PE, and PS phospholipid classes in MDA-MB-231 cells following ACC1 deletion (Figure 3F-H). In MCF7 cells, the effect of ACC1 KO on saturation status was not as consistent in these classes (Figure 3I-K), but the overall effect was that ACC1 deletion shifted the lipidome to a more unsaturated profile in both MCF7 and MDA-MB-231 cells.

**Figure 3.**
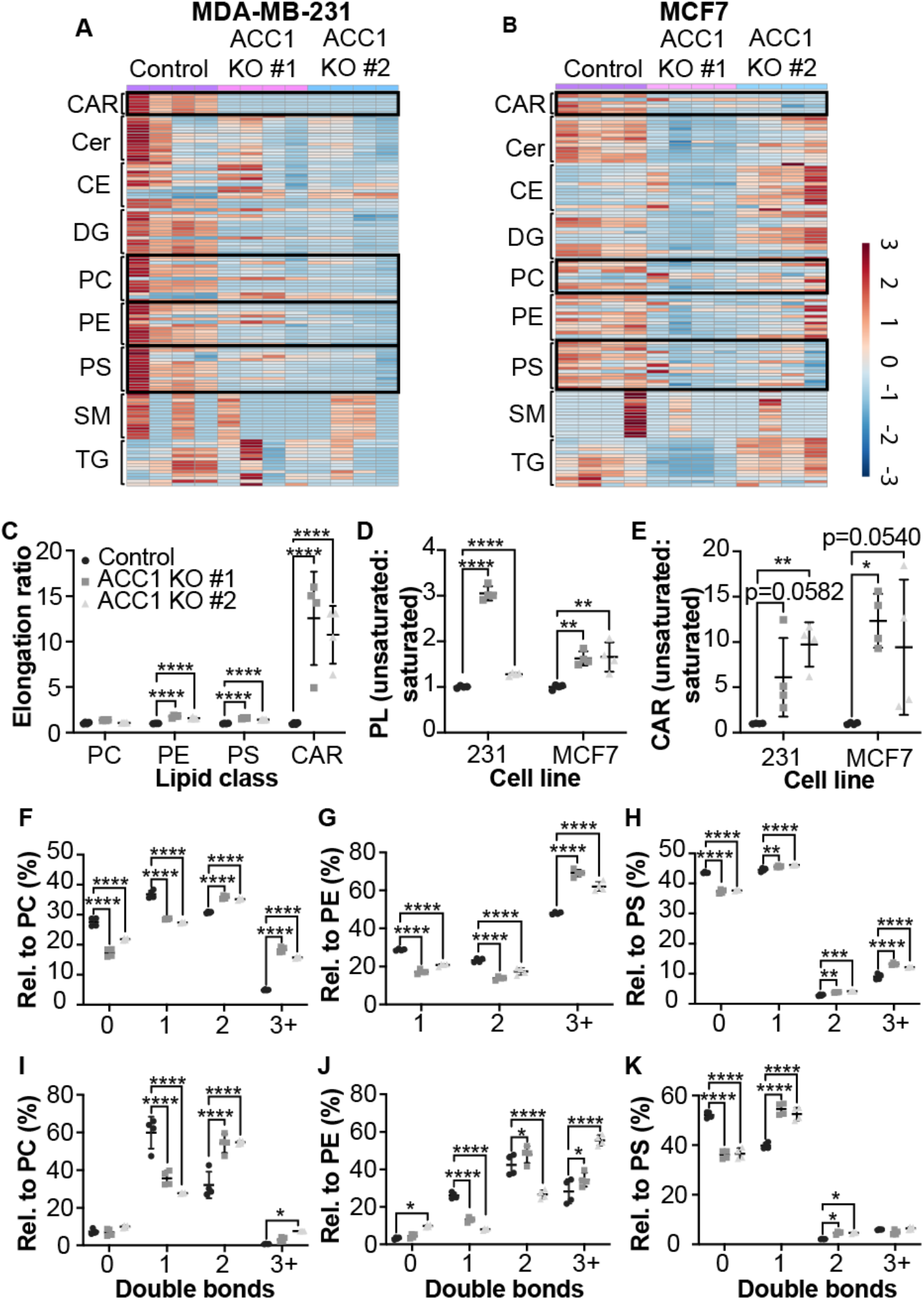
Deletion of ACC1 alters lipid profiles of MDA-MB-231 and MCF7 cells. **(A)** Heat map of 126 lipids in control and ACC1 KO MDA-MB-231 cells from nine lipid classes; acyl-carnitines (CAR), ceramides (Cer), cholesteryl esters (CE), diacylglycerols (DG), phosphatidylcholine (PC), phosphatidylethanolamine (PE), phosphatidylserine (PS), sphingomyelin (SM) and triacylglycerol (TG). **(B)** Heat map of 122 lipids in control and ACC1 KO MCF7 cells from nine lipid classes (as above). Each coloured cell represents lipid abundance normalised to the internal standards of each separate sample with data scaled to the range of each variable (n=4 for each group). Scale ranges from −3 to 3 representing the normalised abundance in arbitrary units. **(C)** The ratio of longer chain:shorter chain PE, PS and CAR in control and ACC1 KO MDA-MB-231 cells. **(D)** Ratio of unsaturated to saturated phospholipids (with control normalised to 1) in control and ACC1 KO MDA-MB-231 and MCF7 cells. **(E)** Ratio of unsaturated to saturated CARs (with control normalised to 1) in control and ACC1 KO MDA-MB-231 and MCF7 cells. Lipid unsaturation profiles in control and ACC1 KO MDA-MB-231 cells of **(F)** PC, **(G)** PE and **(H)** PS represented as a ratio of the total phospholipids in each headgroup (%). Lipid unsaturation profiles in control and ACC1 KO MCF7 cells of **(I)** PC, **(J)** PE and **(K)** PS represented as a ratio of the total phospholipids in each headgroup (%). Data is represented as the mean ± SD (n=4 independent experiments). Statistical significance was determined by an ordinary one-way or two-way ANOVA with Dunnett’s multiple comparisons post hoc test. *P≤0.05, **P≤0.01, ***P≤0.001, ****P≤0.0001.

### Cell morphology and cell number rescued with the addition of palmitic acid

We identified that ACC1 deletion impairs breast cancer progression in vivo, influences breast cancer cell morphology and survival, and results in altered lipid profiles, highlighting the potential importance of DNL in breast cancer cell function. To confirm that ACC1 mediated changes were due to a reduction in *de novo* synthesised lipids, we cultured ACC1 KO cells in cFBS supplemented with palmitate. Palmitate is the initial product of DNL, which is then activated into palmitoyl-CoA, where it can be a substrate for phospholipid synthesis and elongation and/or desaturation reactions. MDA-MB-231 ACC1 KO cells cultured in media supplemented with palmitate did not alter cell morphology given that ACC1 deletion had no effect on the already spindle-like cells (Figure 4A). However, addition of palmitate rescued the decreased live cell number that followed ACC1 deletion to ∼70% of control cells (KO 1: p=0.0606; KO 2: p=0.0135) (Figure 4B). This was accompanied by a decrease in trypan blue positive ACC1 KO cells when compared to the untreated cells (KO 1: p=0.0008; KO 2: p=0.0002) (Figure 4C). On the contrary, addition of palmitate to MCF7 ACC1 KO cells rescued the cell morphology defects in these cells, such that cells re-gained their ability to form tight clusters and lost their spindle-like shape (Figure 4D). Interestingly, the rescue of MCF7 ACC1 KO cell morphology was not associated with a significant increase in the live cell number count (Figure 4E) but resulted in a decrease in trypan blue-positive cells (KO 1: p=0.0258; KO 2: p=0.0055) (Figure 4F).

**Figure 4.**
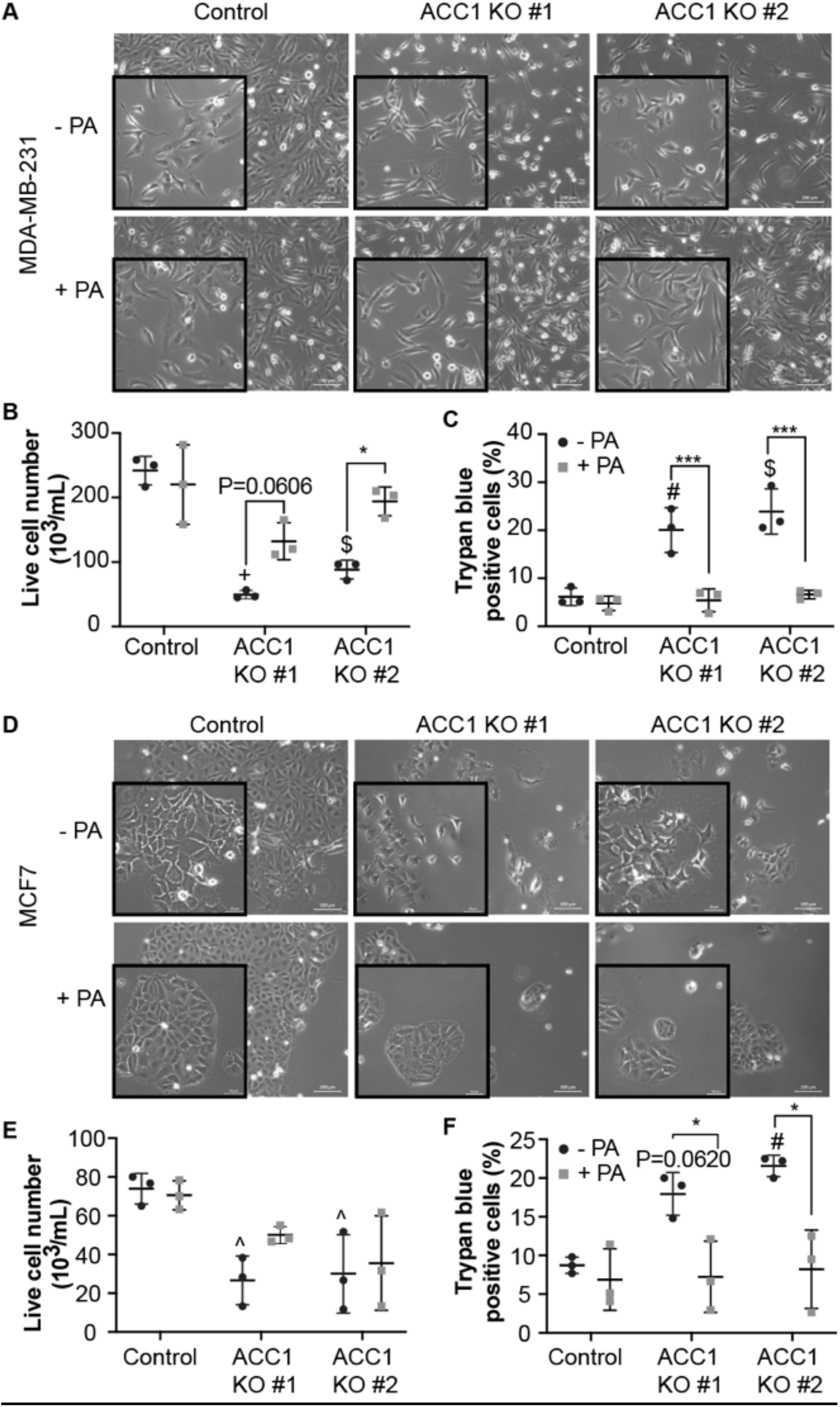
Addition of palmitic acid partially rescues ACC1 KO phenotypes. **(A)** Representative phase contrast images of control and ACC1 KO MDA-MB-231 cells in the presence and absence of exogenous palmitate (PA) 168 h after addition. **(B)** Live and **(C)** trypan blue-positive cell count of MDA-MB-231 control and ACC1 KO cells. **(D)** Representative phase contrast images of control and ACC1 KO MCF7 cells in the presence and absence of exogenous palmitate 168 h after addition. **(E)** Live and **(F)** trypan blue-positive cell count of MCF7 control and ACC1 KO cells. Scale bar = 100μm. Data is represented as the mean ± SEM (n=3 independent experiments). Statistical significance was determined by a 2way ANOVA with Tukey’s multiple comparisons post-hoc test. ^ P≤0.05 #P≤0.01, $P≤0.001, +P≤0.0001 compared to -PA of control; *P≤0.05, ***P≤0.001 compared to -PA for each group.

### ACC1 expression is upregulated in malignant tissue compared to normal adjacent tissue

To investigate the clinical relevance of ACC1 in breast cancer, we measured ACC1 levels in human tissue microarrays (TMAs) containing a selection of breast carcinomas and normal adjacent tissue (NAT). As varied levels of ACC1 staining occurred across the TMA, a scoring index was created based on staining intensity to score all samples (Figure 5A). Normal adjacent tissue was either negative or had very weak ACC1 staining, while malignant tissue expressed higher levels (Figure 5B). To further explore the role of ACC1 in the progression of breast cancer, we compared the ACC1 IHC score of NAT with cancerous cores based on histological grade (Grade 1-3) and stage (Stage I-IIIB). Analysis of ACC1 expression indicated that while all cancerous tissue expressed significantly higher levels of ACC1 compared to NAT, ACC1 levels did not differentiate between higher breast cancer grades or stages (Figure 5C-D). Examination of ACC1 levels in metastatic tissue demonstrated that ACC1 levels are maintained throughout breast cancer progression (Supplementary Figure S6A-E). This data suggests that ACC1 is not expressed or expressed at very low levels in normal breast tissue, with expression increasing during breast cancer initiation which is maintained throughout disease progression, making it an attractive therapeutic target.

**Figure 5.**
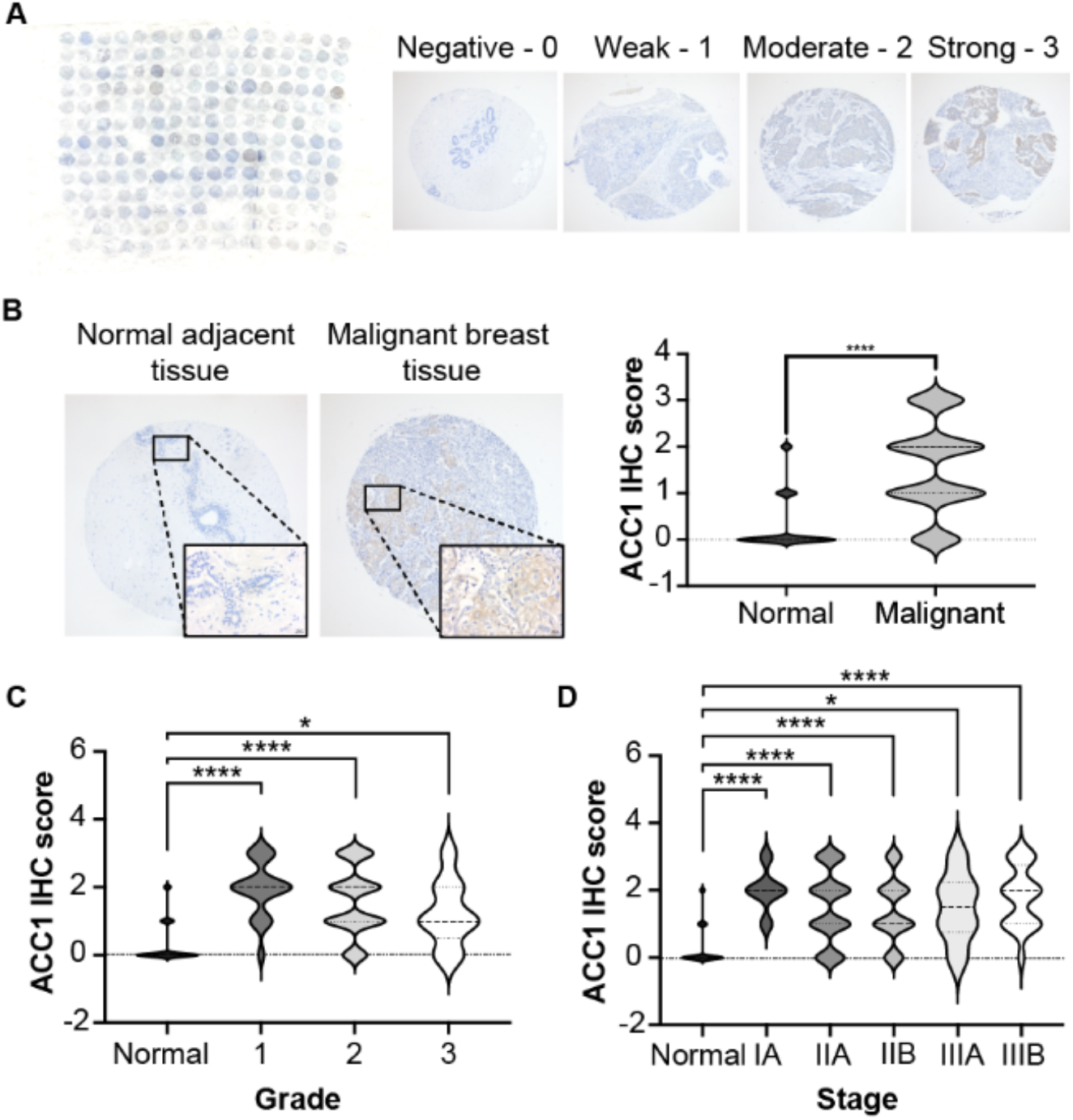
ACC1 expression is upregulated in human breast cancer tissue. **(A)** Representative image of the human breast cancer tissue microarray (TMA) (BR2085d, US Biomax) IHC and the score index used to quantify expression levels of ACC1. **(B)** Representative images of ACC1 immunohistochemistry on normal adjacent and malignant tissue. Scoring of ACC1 expression levels in normal adjacent tissue and the corresponding primary tumours. **(C)** Distribution of ACC1 IHC scores at each tumour histological grade. **(D)** Distribution of ACC1 IHC scores at each tumour stage. Scale bar = 20μm. Data is represented as the mean ± SD (n=38 normal and 167 malignant human cores). Statistical significance was determined by an unpaired student’s t-test or an ordinary one-way ANOVA with Tukey’s multiple comparisons post-hoc text. *P≤0.05, ****P≤0.0001.

## Discussion

This is the first study to demonstrate a critical role for ACC1 in breast cancer progression using mammary specific conditional ACC1 knockout mice and ACC1 CRISPER-Cas9 knockout cancer cell models. Using these models, we have identified a role for ACC1 in driving breast cancer cell proliferation, survival, and membrane function through its role in lipid synthesis, particularly the synthesis of phospholipids and production of acyl-carnitines. Additionally, ACC1 was shown to be a potentially attractive therapeutic target given its low expression profile in normal breast and high expression during breast cancer progression and metastasis.

The role of ACC1 in a physiological metabolic context is well understood [29–32] but its role in cancer, specifically breast cancer, remains unclear. We used genetic knockout of ACC1 in two different cell lines to demonstrate an *in vitro* role for the enzyme in cancer cell growth and survival. Several other investigators have used transient genetic and pharmacological approaches against ACC1 to demonstrate a potential role in breast cancer cell proliferation, survival and metastasis following treatment [9, 10, 24] with others demonstrating a role for ACC in other cancer types [33, 34]. Expanding on the *in vitro* findings of ours and others, we describe for the first time, the *in vivo* effects of BlgCre-promoter driven ACC1 deletion in a PyMT mammary tumour mouse model. Deletion of ACC1 resulted in an increase in tumour-free and overall survival, which was driven by a decrease in cancer cell proliferation. A recent publication by Marczyk and colleagues complements this study by demonstrating that pharmacological inhibition of both ACC1/ACC2 resulted in decreased tumour growth in mouse MDA-MB-468 xenografts and patient-derived TNBC xenografts [23]. Together these studies indicate the translational potential of targeting ACC1 and the importance of this enzyme in breast cancer.

Extending upon the study by Stoiber and colleagues [9], who demonstrated that treatment of SKBR3 breast cancer cells with Soraphen A resulted in altered membrane mechanics, we characterised cell morphological changes of MCF7 cells following genetic deletion of ACC1. Morphological changes encompass alterations in cell shape, structure and size, which can determine the survival of cells [35]. We show that ACC1 deletion altered all three aspects of cell morphology, highlighting the importance of ACC1-derived lipids in the formation of cell membranes and membrane structures. It is well established that phospholipids represent the major lipid component of cell membranes, and a few studies have begun to link the alteration of phospholipid composition to these morphological changes, which ultimately contributes to the observed phenotypes in cancer [9, 36, 37]. Our study demonstrates that deletion of ACC1 decreases levels of PC, PE, PS species, while also shifting the lipidome to more elongated and less saturated lipids. Changes in the characteristics of membrane lipids in addition to the overall reduction in phospholipids likely underpin the observed decreased proliferation. In our study, addition of non-toxic concentrations of palmitate, a saturated FA, to an environment of elevated MUFAs and PUFAs, decreased cell death. These data add to the growing evidence that the balance and relationship between SFA, MUFA and PUFAs are important in maintaining cell integrity and survival, and we highlight an important role for ACC1 maintaining this homeostasis [38–40]. As the rescue was only partially effective in MCF7 cells, this may suggest that ACC1 has other roles in breast cancer cells beyond its documented lipogenic role.

Lipid integrity and composition are important for cell-cell adhesion [41] and cell-matrix adhesion [42]. Here, we show that deletion of ACC1 alters MCF7 cell shape and structure due to modification of phospholipid composition. Cells displayed more spindle-like processes and a decreased ability to form cell-cell adhesions in 2D adherent culture, as well as redistribution of F-actin resulting in a distinct increase in actin rich peripheral ruffles. Membrane ruffles are actin rich processes formed from lamellipodia that failed to attach to the substratum, resulting in delayed and decreased cell-matrix adhesion [43–46]. Cell-cell and cell-matrix adhesions are mediated by cadherins and integrins, respectively [47, 48], and are critical in the process of collective migration, proliferation and survival [49, 50], thus interference of these processes through abrogating ACC1 may provide a promising therapeutic target for breast cancer.

Interestingly, deletion of ACC1 increased MCF7 cell size, suggesting a potential disruption in the balance between growth and proliferation [51, 52], as the absence of ACC1 was also associated with decreased proliferation. A study by Miettinen and colleagues [52] identified that lipid levels and cell size are negatively correlated, such that larger cells are associated with decreased lipid availability, which is evident in our study. As well as lipid availability, mitochondrial function is also a factor contributing to maintaining cell size homeostasis [52]. Although we did not measure mitochondrial function directly, the abundance of CAR species was significantly lower in ACC1 KO cells, which can be reflective of mitochondrial ß-oxidation status [50]. Further characterisation of the CAR species may elucidate the significance of the decrease following ACC1 deletion.

ACC1 levels in human tissue has previously been explored, with Pandey and colleagues [53] reporting that ACC1 expression was almost undetectable in normal adjacent breast tissue but upregulated in ductal carcinoma *in situ* (DCIS). DCIS is the most common type of non-invasive breast cancer which has the potential to become invasive, thus represents a step between normal breast tissue and invasive breast cancer [54]. Our study further observed ACC1 expression in invasive breast cancer, when compared to normal adjacent breast tissue. Together, these data indicate that ACC1 is highly expressed both in localised and invasive breast tumours, but may also have a role in establishing disseminated cells at distal sites, given that ACC1 remains highly expressed in distal metastatic tissue. Overall, this data highlights the importance of ACC1 in the initiation of breast cancer and progression of the primary tumour, given the presence and increased expression of ACC1 at early and late stages of breast cancer, when compared to normal human breast tissue.

In summary, we utilised two genetic deletion models to investigate the impact of ACC1 in breast cancer. We demonstrated a role for ACC1 in driving progression of breast cancer through its role in proliferation and its impact on lipid profiles, while analysis of human breast tissue demonstrated upregulation of ACC1 in malignant tissue compared to normal tissue. We believe this work indicates that targeting ACC1 may have therapeutic potential for the treatment of breast cancer.

## Additional information

This work was supported by Project Grant 1100626 and Ideas Grant 2002660 from the National Health and Medical Research Council of Australia (A.S.D) and Cancer Council NSW Research Project Grant (RG 20-08) and Priority-driven Collaborative Cancer Research Scheme (Grant #1130499), funded by the National Breast Cancer Foundation Australia with the assistance of Cancer Australia (M.J.N).

## Conflicts of Interest

The authors declare that the research was conducted in the absence of any commercial or financial relationships that could be construed as a potential conflict of interest.

## Notes

### Competing Interest Statement

The authors have declared no competing interest.

