## Supplemental text file for "Acetyl-CoA carboxylase 1-dependent lipogenesis drives breast cancer progression"

**Supplementary data**

**Supplementary Table S1. List of internal standards used in mass spectrometry analysis.**

Names and final amounts (nmol) of the lipids used in the internal standard mixture for normalisation, per sample. All standards were purchased from Avanti Polar Lipids, USA or Cayman Chemical, USA.

| Lipid name | Final amount (nmol) |
| --- | --- |
| 17:0 sphingosine | 0.2 |
| 17:0 sphingosine-1-phosphate | 0.2 |
| 18:1-d7 monoacylglycerol | 0.2 |
| 18:1/12:0 lactosylceramide | 0.2 |
| Palmitoyl-L-carnitine-(N-methyl-d3) | 0.5 |
| 18:1/15:0-d7 diacylglycerol | 0.5 |
| 17:0 lysophosphatidylcholine | 0.5 |
| 17:1 LPE | 0.5 |
| 18:1/17:0 ceramide | 1.0 |
| 17:01/17:0 phosphatidylglycerol | 1.0 |
| 14:0/14:0/14:0/14:0 cardiolipin | 1.0 |
| 17:0/17:0 phosphatidylserine | 1.0 |
| 17:0/17:0 phosphatidylethanolamine | 1.0 |
| 17:0/17:0 phosphatidic acid | 1.0 |
| 17:0/17:0/17:0 triacylglycerol | 1.0 |
| 18:1/12:0 sphingomyelin | 1.0 |
| 18:1/12:0 glucosylceramide | 1.0 |
| 18:1/15:0-d7 phosphatidylinositol | 1.2 |
| 17:0 cholesteryl ester | 2.0 |
| 19:0/19:0 phosphatidylcholine | 2.0 |

**Supplementary Figure S1. Deletion of ACC1 has no effect of mammosphere number or size.**

**(A)** Number of mammospheres formed from control and ACC1 knockout MDA-MB-231 and MCF7 cells. Diameter of mammospheres represented as a % of the total number of mammospheres formed from control and ACC1 KO **(B)** MDA-MB-231 and **(C)** MCF7 cells. Data is represented as the mean ± SEM (n=3 independent experiments). Statistical significance was determined by a two-way ANOVA with Dunnett’s multiple comparisons post-hoc test. ***P≤0.001, ****P≤0.0001.

**Supplementary Figure S2. ACC1 KO has no impact on cell morphology in MDA-MB-231 cells.**

**(A)** Quantification of MDAMB-231 cell size using ImageJ following ACC1 KO **(B)** Quantification of membrane ruffling using ImageJ software in MDA-MB-231 cells. Data is represented as the mean ± SEM (n=3 independent experiments). Statistical significance was determined by an ordinary one-way ANOVA with Dunnett’s multiple comparisons post hoc test. *P≤0.05.

**Supplementary Figure S3. Deletion of ACC1 decreases total lipid levels of PC, PE, PS, and CAR classes in MDA-MB-231 cells.**

**(A)** Total lipid levels of PC, PE, PS, and CAR in MDA-MB-231 control and ACC1 KO cells. Data is represented as the mean ± SD (n=4 independent experiments). Statistical significance was determined by an ordinary one-way ANOVA with Dunnett’s multiple comparisons post hoc test. **P≤0.01, ***P≤0.001.

**Supplementary Figure S4. Deletion of ACC1 decreases total lipid levels of PC, PS, and CAR classes in MCF7 cells.**

**(A)** Total lipid levels of PC, PS, and CAR in MCF7 control and ACC1 KO cells. Data is represented as the mean ± SD (n=4 independent experiments). Statistical significance was determined by an ordinary one-way ANOVA with Dunnett’s multiple comparisons post hoc test. *P≤0.05, **P≤0.01, ***P≤0.001, ****P≤0.0001.

**Supplementary Figure S5. Deletion of ACC1 has no effect on lipid elongation in MCF7 cells.**

The ratio of longer chains:shorter chains of PE, PS and CARs in control and ACC1 KO MCF7 cells. Longer PL chains were defined as ≥ 36 carbons in the two acyl chains combined and shorter chains were defined as ≤ 35 carbons in the two acyl chains combined. For CARs, longer chains were defined as ≥ 18 carbons and shorter chains defined as ≤17 carbons lipids.
**Supplementary Figure S6. Primary breast tumour and corresponding metastatic lymph node have similar expression levels of ACC1.**

**(A)** Representative image of the human breast cancer tissue microarray (TMA (BR10010b, US Biomax) IHC and the score index used to quantify expression levels of ACC1. **(B)** Correlation of ACC1 IHC scoring index with % of PR staining (R=-0.2292, R2=0.0525). **(C)** Correlation of ACC1 IHC scoring index with % of ER staining (R=-0.0290, R2=0.0008). **(D)** Correlation of ACC1 IHC scoring index with HER2+ IHC scoring index (R=0.2468, R2=0.0609). **(E)** Scoring of ACC1 expression levels in primary tumours and the corresponding metastatic lymph nodes. **(**Scale bar = 20μm. Data is represented as the mean ± SD (n=50 tumour and 50 corresponding metastatic lymph node human cores). Statistical significance was determined by an unpaired student’s t-test and a Pearson’s correlation.
