## Supplementary figures and images for "Acetyl-CoA carboxylase 1-dependent lipogenesis drives breast cancer progression"

### Supplemental Figure S1

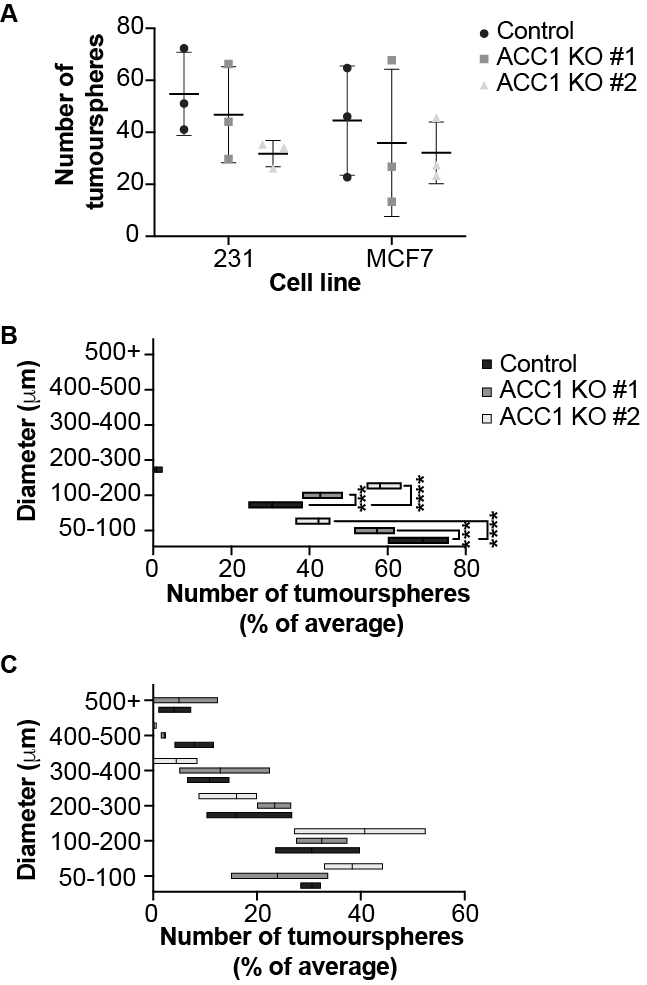

### Supplemental Figure S2

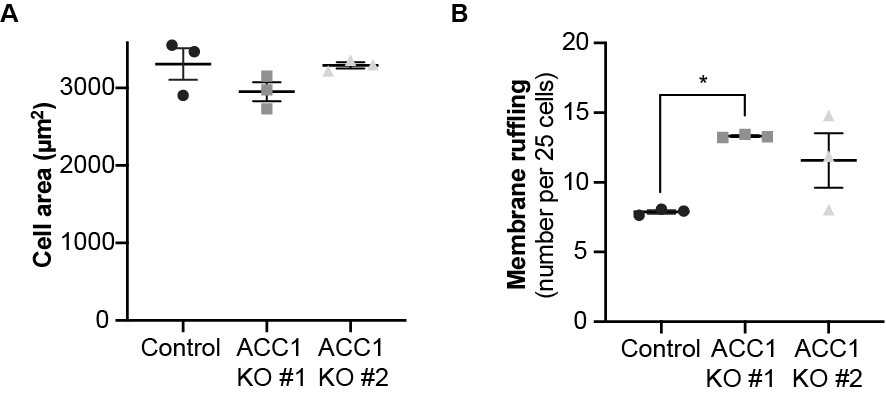

### Supplemental Figure S3

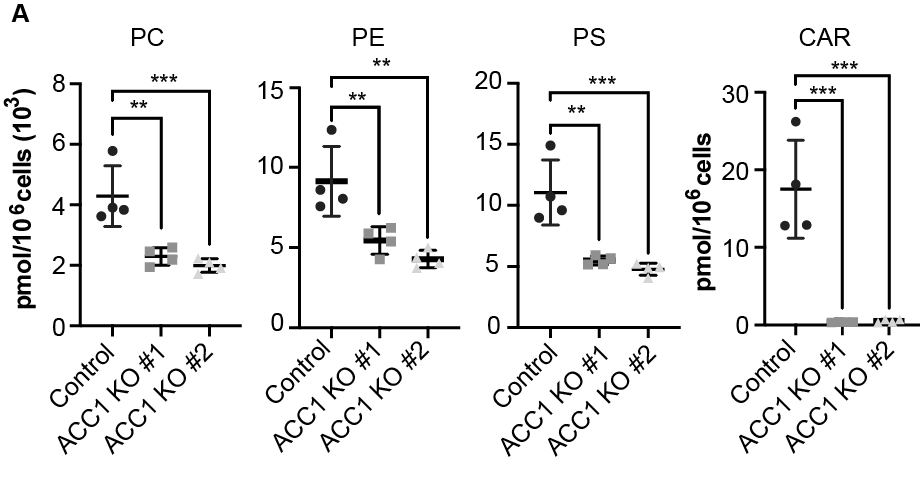

### Supplemental Figure S4

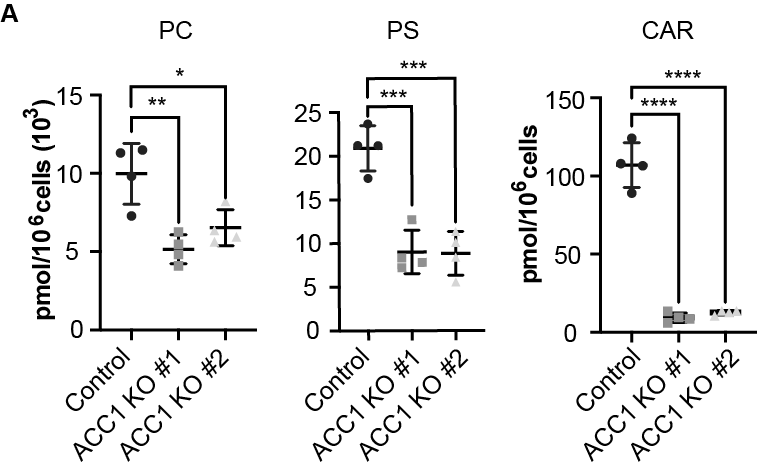

### Supplemental Figure S5

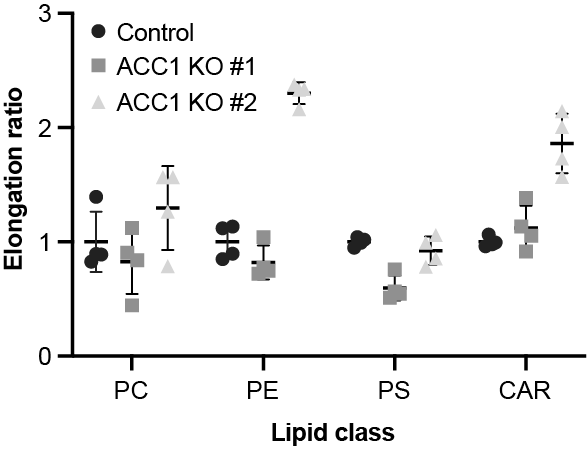

### Supplemental Figure S6

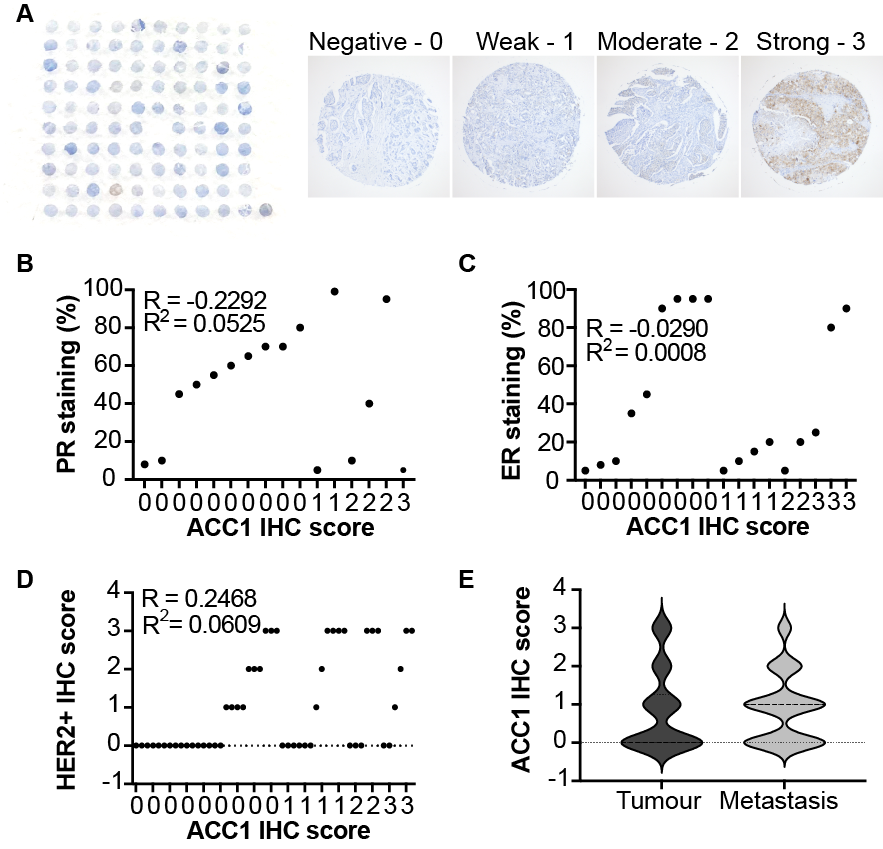
